# MSF-CPMP: A Novel Multi-Source Feature Fusion Model for Cyclic Peptide Membrane Permeability Prediction

**DOI:** 10.1101/2025.07.28.667130

**Authors:** Yijun Zhang, Zimeng Chen, Zhuxuan Wan, Qianhui Jiang, Xiaoling Lu, Bin Yan, Jing Qin, Yong Liu, Junwen Wang

**Affiliations:** School of Physics and Electronic Information, Guangxi University for Nationalities, Nanning, Guangxi Province, China; School of Artificial Intelligence, Guangxi University for Nationalities, Nanning, Guangxi Province, China; Division of Applied Oral Sciences & Community Dental Care, Faculty of Dentistry, The University of Hong Kong, Hong Kong SAR, China; State Key Laboratory of Pharmaceutical Biotechnology, The University of Hong Kong, Hong Kong SAR, China; School of Pharmaceutical Sciences (Shenzhen), Shenzhen Campus of Sun Yat-sen University, Shenzhen, Guangdong Province, China

## Abstract

Cyclic peptides are becoming attractive molecules for drug discovery because of their properties with inherent stability and structural diversity. However, the high potential of cyclic peptide drugs is challenged by the limited membrane permeability cross cell membrane. To predict cyclic peptide membrane permeability (CPMP), an increased number of computational models or tools are designed and used. But these existing algorithms or models do not appropriately capture feature diversity of cyclic peptides. In this study, we introduce a novel multi-source feature fusion model called MSF-CPMP, which aims to increase the accuracy of predicted CPMP. The MSF-CPMP model incorporates three features extracted from SMILES sequences, graph-based molecular structures, and physicochemical properties of cyclic peptides. By benchmarking with other machine learning and deep learning-based methods, MSF-CPMP achieved the highest levels of the evaluation metrics such as accuracy of 0.9062 and AUROC of 0.9546, and further validated MSF-CPMP robustness in learning capabilities and efficacy of its multi-source fusion. Our result demonstrates that MSF-CPMP outperforms other methods in predicting CPMP, that provides also exemplifies the power of advanced deep learning methods in tackling complex biological challenges, offering contributions to computational biology and clinical treatment. Code is available at https://github.com/wanglabhku/MSF-CPMP

**Author summary:** We have recognized that cyclic peptides are one type of main macro-molecular drugs useful for treatment of human diseases. However, the structural diversity of cyclic peptides results in limited permeability across cell membranes which is challenging drug research or industrial development. Previously some computational models or tools including machine learning or deep learning ones tried to solve this problem, but they still are not efficient. There are urgent requirement for more effective methodologies. Thus, we introduce a novel multi-source feature fusion model called MSF-CPMP. Our aim is to enhance accuracy and efficiency of predicted CPMP. We carried out evaluation of performance metrics by comparative analysis between MSF-CPMP and other machine learning or deep learning methods. Our result demonstrates that MSF-CPMP outperforms other models in predicting CPMP. We further validated MSF-CPMP robustness in learning capabilities and efficacy of its multi-source feature fusion. In the future we would further improve our methods that can integrate broader biomedicine and biomedical information and provide guide for drug diversity in clinically treating complex human diseases.

## Introduction

Cyclic peptides are attractive in drug development because of their closed-loop structure that can offer exceptional stability, bioactivity, and resistance to enzymatic degradation. These values of cyclic peptides have let them become important in drug discovery [1–4]. Membrane permeability refers to ability of drugs to cross cell membranes and is one of main factors determining whether a drug is successfully absorbed into target sites in human body. With optimal level of permeability is mainly considered, however, limited permeability of cyclic peptides, especially for oral administration, hinders largely clinical application of this kind of drugs [5–7]. Such lower membrane permeability directly affects the absorption, distribution, metabolism, and excretion profiles of cyclic peptides, and thereby results in a decreased therapeutic efficiency. Thus, selection of higher cyclic peptide membrane permeability (CPMP) is a major goal during early stage of drugs treating human disease, particularly complex diseases, such as cancers, where effective tumor cell penetration is vital [8–10].

Traditional approaches to measure CPMP rely mainly on labor-intensive experimental methods, such as Caco-2 cell assays, which are costly and time-consuming [11–13]. Early studies using computational methods, like those by Patric et al. and Taha et al., established foundational models that can correlate molecular properties with permeability but lack scalability [14, 15]. Recent advancements have introduced deep learning algorithms, such as the CycPeptMP model, that designs cyclic peptide features at atomic, monomer, and peptide levels, achieves a mean absolute error of 0.355 and a correlation coefficient of 0.883 using deep learning techniques [16]. Cao et al. introduced Multi CycGT model, a multimodal model combining a Graph Convolutional Network (GCN) and a transformer to extract one- and two-dimensional features for predicting CPMP, achieving an accuracy (ACC) of 0.8206 [17]. Despite partially effective, these reported models often fail to fully capture the intricate structural and physical/chemical features of cyclic peptides, thus still limiting their power to identify the optimal rates of CPMP [18].

To accelerate cyclic peptide drug discovery, more efficient computational methods or tools that perform most likely accurate prediction of CPMP are urgently required. Recently, deep learning methods extracting multi-dimensional features from complex peptide data, are expected to offer innovative solutions that leverage advanced machine learning techniques [19, 20]. In this study, we propose MSF-CPMP, a novel model that can integrate multi-source features extracted from SMILES sequences, graph-based molecular structures, and physicochemical properties. By employing transformer-based natural language processing for sequence analysis and graph neural networks for structural feature learning, our method captures both local and global molecular characteristics. Extensive benchmarking against eight machine learning and ten deep learning models demonstrates the superiority of MSF-CPMP in predicting CPMP. Feature visualization and ablation testing further validate the robustness of MSF-CPMP in learning capabilities and the efficacy of its multi-source integration. MSF-CPMP exemplifies the power of advanced deep learning methods in tackling complex biological challenges, offering significant contributions to computational biology and drug discovery.

## Results

### Overview of MSF-CPMP framework

Fig 1 displays the schematic diagram of MSF-CPMP modelling system that integrates multi-source features and aims to predict membrane permeability cyclic peptides (CPMP). The MSF-CPMP framework consisting of three steps. In data pre-processing (Fig 1), MSF-CPMP employs a Siamese network with two parallel branches, each containing transformer encoder layers, with shared weight parameters. This similarities between input features while modelling long-range dependencies in the SMILES sequences.

**Fig 1.**
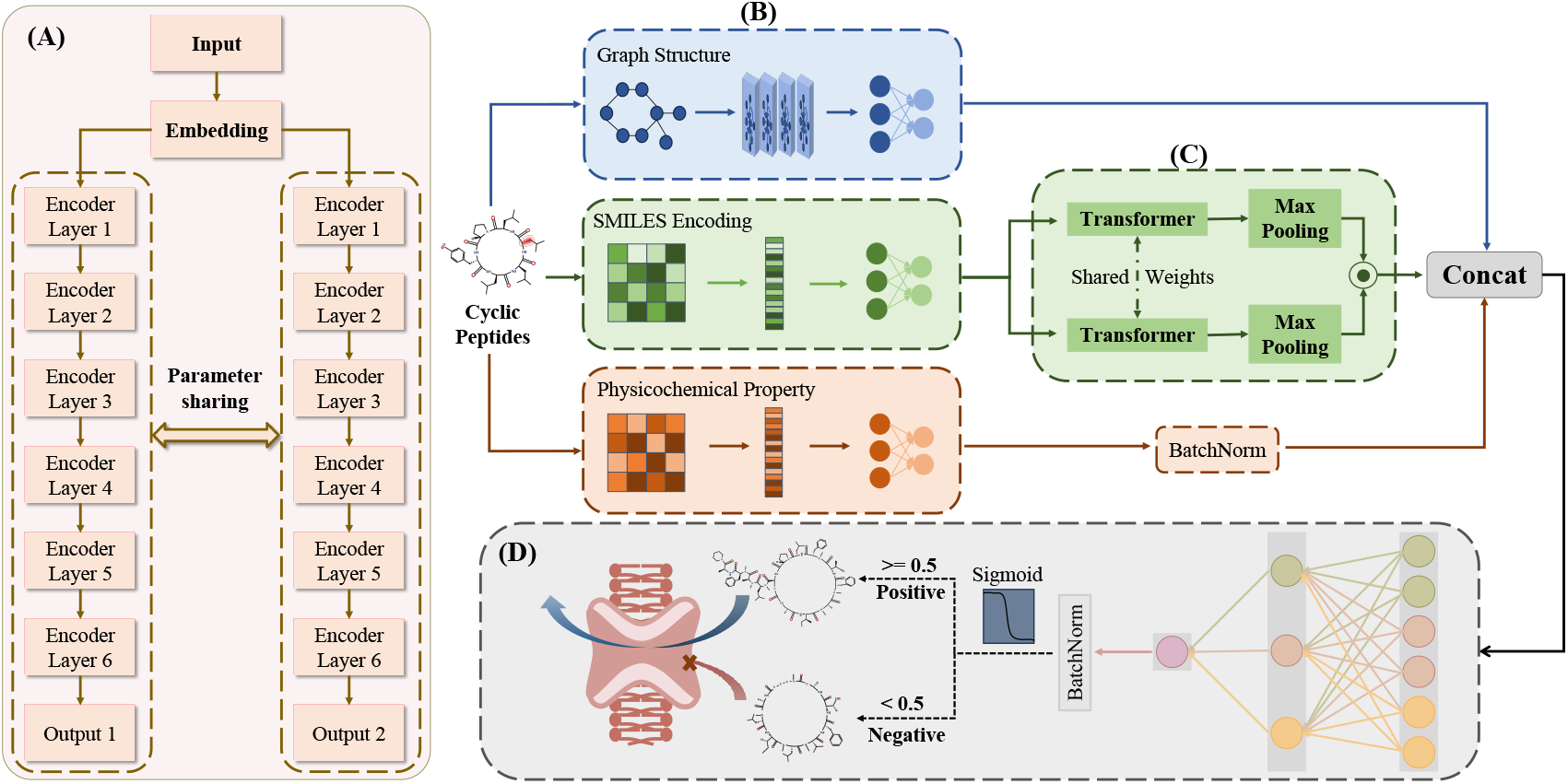
Schematic diagram of the MSF-CPMP model. (A) The MSF-CPMP model employs a Siamese network with two parallel branches, each containing 6 Transformer encoder layers with shared weight parameters, capturing similarities between input features while modeling long-range dependencies in the SMILES sequences. (B) The MSF-CPMP model considers the graph structure, SMILES encoding, and physicochemical properties of each cyclic peptide. First, a graph attention network is used to embed the graph structure, and two artificial neural networks are used to embed the SMILES sequence and physicochemical properties. (C) The Siamese network uses two Transformer modules to embed the processed SMILES sequence and then performs a Hadamard product. (D) The MSF-CPMP model then uses a feature fusion module to extract important cyclic peptide feature information and predict the membrane permeability of the cyclic peptide.

1. Firstly, the MSF-CPMP system considers the graph structure, SMILES encoding, and physicochemical properties of each cyclic peptide. A graph attention network is used to embed the graph structure, and two artificial neural networks are used to embed the SMILES sequence and physicochemical properties.
2. Secondly, the Siamese network uses two Transformer modules to embed the processed SMILES sequence and then performs a Hadamard product.
3. Thirdly, the MSF-CPMP model then uses a feature fusion module to extract important cyclic peptide feature information and predict the membrane permeability of the cyclic peptides.

MSF-CPMP is a deep learning and multi-source feature fusion model. It combines, three distinct feature representations of cyclic peptides were incorporated: graph structures, SMILES string encodings, and physicochemical properties. The detailed tokenization steps and representative examples are illustrated in Figure S3 and Figure S4. The architectural parameters and configurations for each module in the MSF-CPMP model are detailed in Table S1. The input and output dimensions for each module used in MSF-CPMP are enumerated (Table S1).

### Evaluation of MSF-CPMP by comparing with machine learning models

In order to evaluate performance of MSF-CPMP, we first compared with eight traditional machine learning methods. Cyclic peptide datasets are extracted from CycPeptMPDB (Cyclic Peptide Membrane Permeability Database), including 103 physicochemical properties and SMILES string data of cyclic peptides were subject to these models and to predict CPMP. The top ten cyclic peptides predicted by MSF-CPMP are listed (Table S2), all of which are positive. The cyclic peptide PEPTIDE7214 has the highest predicted probability.

Through ten-fold cross-validation, we observed that MSF-CPMP exhibits the highest levels of almost every metrics (Table S3, Table 1). ACC value of MSF-CPMP exceeds other methods by 0.098 ∼ 0.143 in ACC (Table 1). By exhibiting Area Under the Receiver Operating Characteristic curve (AUROC) and Area Under the Precision-Recall curves (AUPRC), MSF-CPMP achieved the largest values Fig 2. Next, we calculated regression metrics to validate performance of MSF-CPMP. We observed lower levels of MSE and MAE, and the highest *R*^2^ (Table 1). A more comprehensive evaluation metrics of classifications (including ACC, Precision, F1 Score, Recall, MCC, TNR, and KAPPA) are provided in Table S3, of regression metrics (MSE, MAE, RMSE, MAPE, *R*^2^ and r) in Table S4. This finding supports for our proposed MSF-CPMP that surpasses other machine learning methods.

**Table 1.** Comparison of MSF-CPMP performance with selected machine learning models.

| Models | ACC | Precision | Recall | MSE | MAE | $R^2$ |
| --- | --- | --- | --- | --- | --- | --- |
| **MSF-CPMP** | **<u>0.9062</u>** | **<u>0.8937</u>** | **<u>0.9916</u>** | **<u>0.0917</u>** | **<u>0.2428</u>** | **<u>0.5774</u>** |
| CatBoost | 0.7791 | 0.8016 | 0.8889 | 0.4018 | 0.505 | 0.1584 |
| XGBoost | <u>0.8067</u> | 0.8379 | 0.8926 | 0.446 | 0.4382 | 0.1056 |
| KNN | 0.7631 | 0.7829 | <u>0.9148</u> | 0.3704 | 0.5774 | -1.2557 |
| LGBM | 0.798 | 0.8136 | 0.9153 | 0.453 | 0.4463 | 0.2732 |
| RF | 0.7951 | 0.8414 | 0.8711 | 0.6349 | 0.4406 | 0.0726 |
| SVM (poly) | 0.7965 | 0.836 | 0.875 | 0.4768 | 0.4837 | -0.3034 |
| SVM (rbf) | 0.8081 | 0.8423 | 0.8884 | <u>0.3087</u> | <u>0.3822</u> | <u>0.2214</u> |
| DT | 0.7863 | <u>0.8495</u> | 0.8369 | 0.0917 | 0.3657 | 0.3146 |

Bold and underlined values represent the best and the second-best performance for each metrics, respectively.

**Fig 2.**
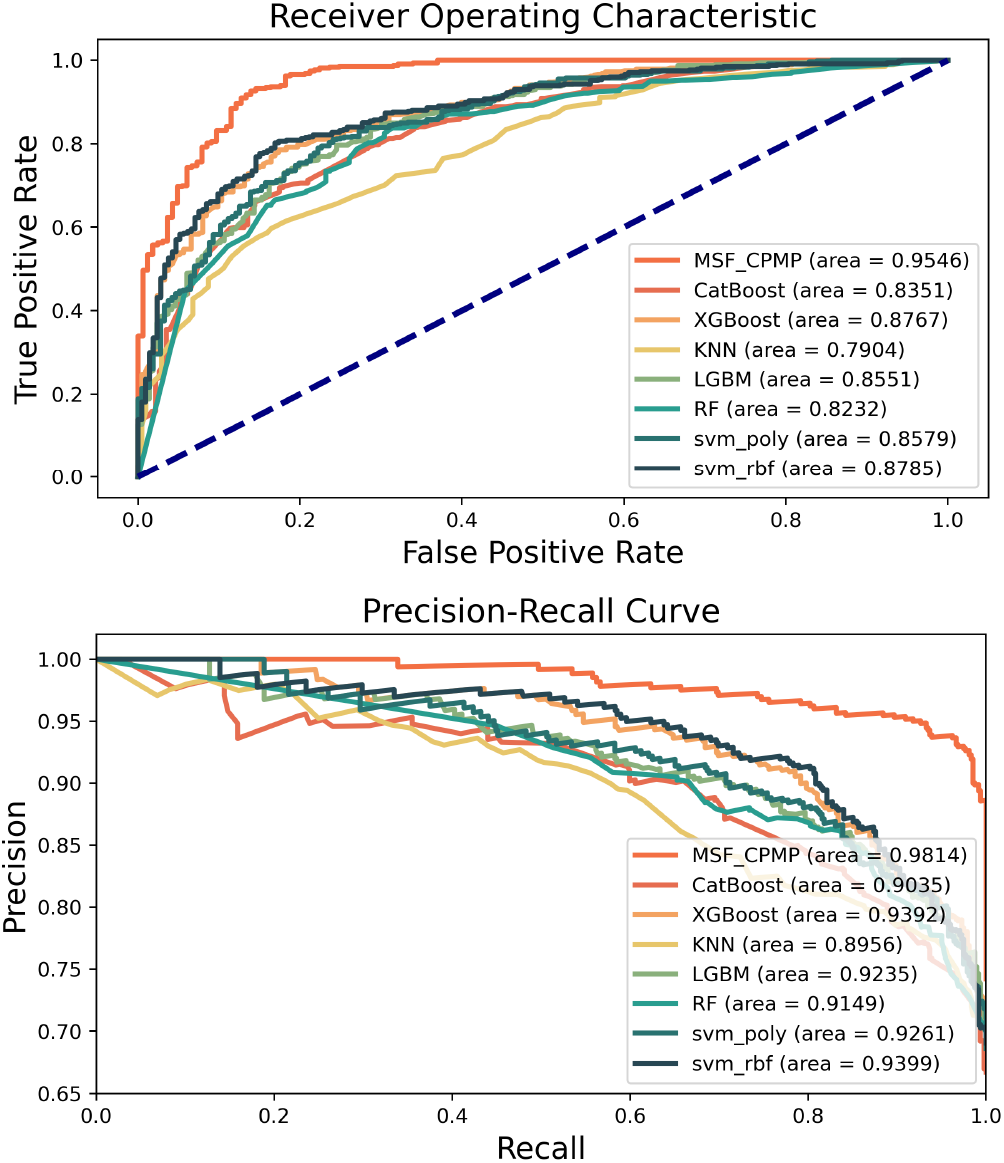
Comparison of predictive performance beween MSF-CPMP and different machine learning methods. (A) Area Under the Receiver Operating Characteristic curve (AUROC) and (B) Area Under the Precision-Recall curves (AUPRC).

### Evaluation of MSF-CPMP by comparing with deep learning models

Deep learning models have been widely used in drug research, such as Multi CycGT, CNN, GAT, GCN, GRU, LSTM, MLP, Multihead, RNN, and Transformer. Here, we contacted a similar comparison between MSF-CPMP and the ten deep learning methods. As a result, deep learning-based MSF-CPMP was found in top position in seven individual metrics (ACC, Precision, F1, KAPPA, MCC, TNR and KAPPA), with only ranked the second of Recall (Table 2, Table S5). Notably, Multi CycGT ranked the second in four metrics (ACC, Precision, F1 and KAPPA). MSF-CPMP also gained the largest values of AUROC and AUPRC (Fig 3) among the deep learning-based models. This finding suggests under consistent input conditions, sigmoid layer removed with smooth L1 loss, that MSF-CPMP exhibits superior stability with the lowest levels of MSE and MAE, and highest value of *R*^2^ (Table 2, Table S6), indicating its minimized variance and enhanced determination in the prediction.

**Table 2.**
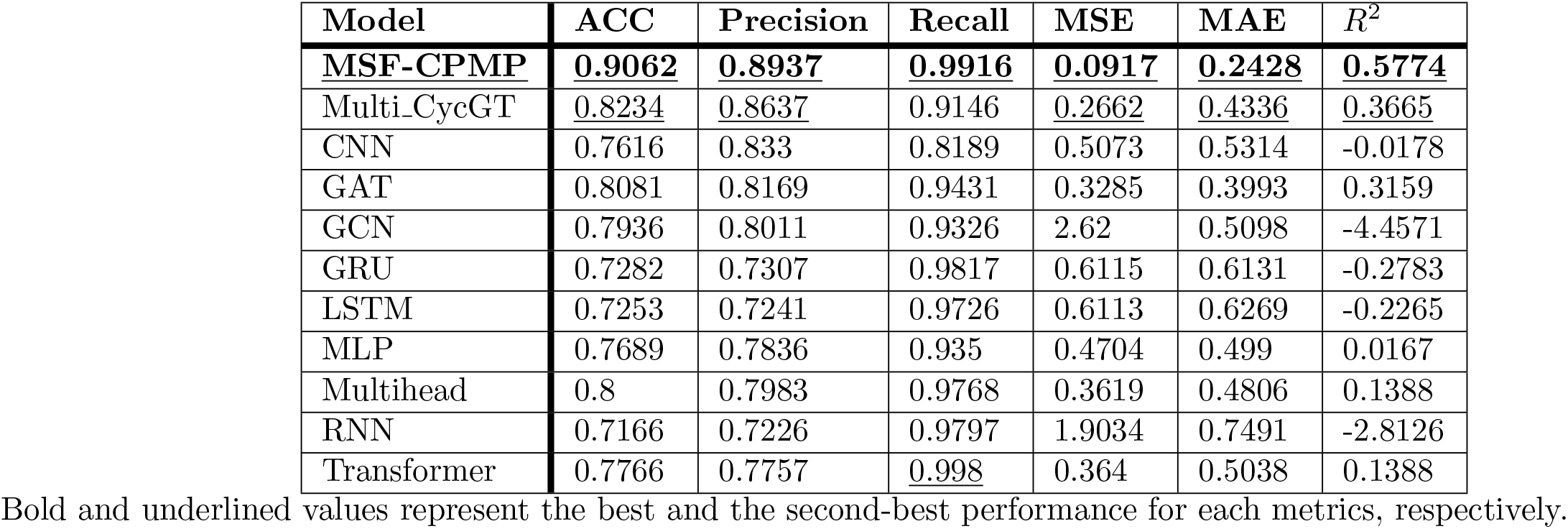
Performance comparison of MSF-CPMP with selected deep learning models.

| Model | ACC | Precision | Recall | MSE | MAE | $R^2$ |
| --- | --- | --- | --- | --- | --- | --- |
| **MSF-CPMP** | **<u>0.9062</u>** | **<u>0.8937</u>** | **<u>0.9916</u>** | **<u>0.0917</u>** | **<u>0.2428</u>** | **<u>0.5774</u>** |
| Multi_CycGT | 0.8234 | 0.8637 | 0.9146 | 0.2662 | 0.4336 | 0.3665 |
| CNN | 0.7616 | 0.833 | 0.8189 | 0.5073 | 0.5314 | -0.0178 |
| GAT | 0.8081 | 0.8169 | 0.9431 | 0.3285 | 0.3993 | 0.3159 |
| GCN | 0.7936 | 0.8011 | 0.9326 | 2.62 | 0.5098 | -4.4571 |
| GRU | 0.7282 | 0.7307 | 0.9817 | 0.6115 | 0.6131 | -0.2783 |
| LSTM | 0.7253 | 0.7241 | 0.9726 | 0.6113 | 0.6269 | -0.2265 |
| MLP | 0.7689 | 0.7836 | 0.935 | 0.4704 | 0.499 | 0.0167 |
| Multihead | 0.8 | 0.7983 | 0.9768 | 0.3619 | 0.4806 | 0.1388 |
| RNN | 0.7166 | 0.7226 | 0.9797 | 1.9034 | 0.7491 | -2.8126 |
| Transformer | 0.7766 | 0.7757 | <u>0.998</u> | 0.364 | 0.5038 | 0.1388 |

Bold and underlined values represent the best and the second-best performance for each metrics, respectively.

**Fig 3.**
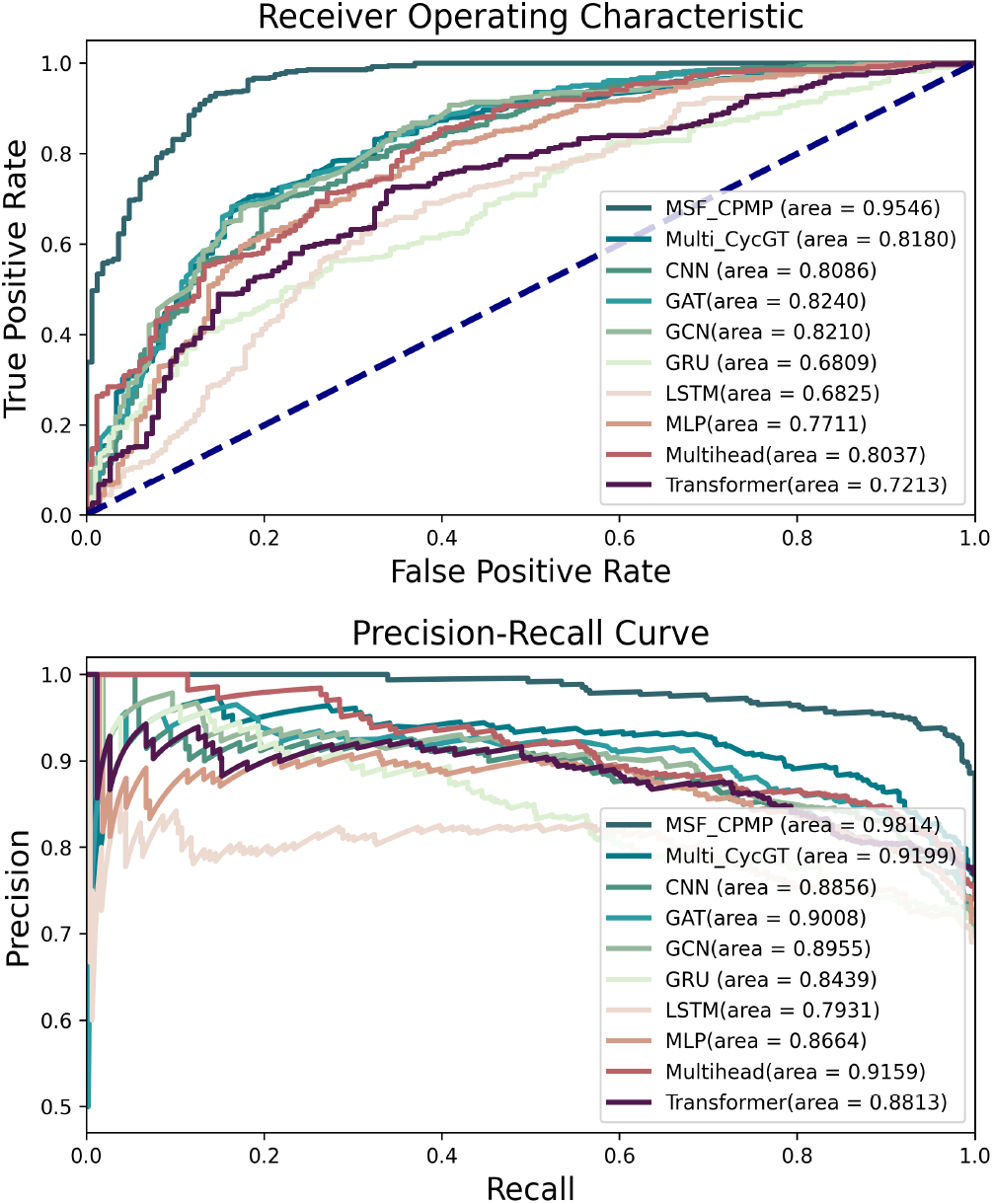
Comparison of predictive performance beween MSF-CPMP and different deep learning methods. (A) Area Under the Receiver Operating Characteristic curve (AUROC) and (B) Area Under the Precision-Recall curves (AUPRC).

### Structural effect of cyclic peptides on MSF-CPMP

To discern subtle structural effects of MSF-CPMP performace, we validated through two critical cases. First, for structurally similar negative cyclic peptides differing only in a single triangular carbon (highlighted in red, Fig 4A1 and A2), MSF-CPMP correctly classified both with low permeability probabilities (e.g., 0.436), revealing its sensitivity to minimal steric perturbations that impede membrane penetration. Second, in a challenging positive sample (Fig 4B1), MSF-CPMP achieved a high probability (0.925) of prediction, whereas Multi CycGT (0.280) and LSTM (0.145) failed to classify it as positive. This result underscores MSF-CPMP ‘s robustness in capturing complex structural determinants overlooked by specialized or sequential models.

**Fig 4.**
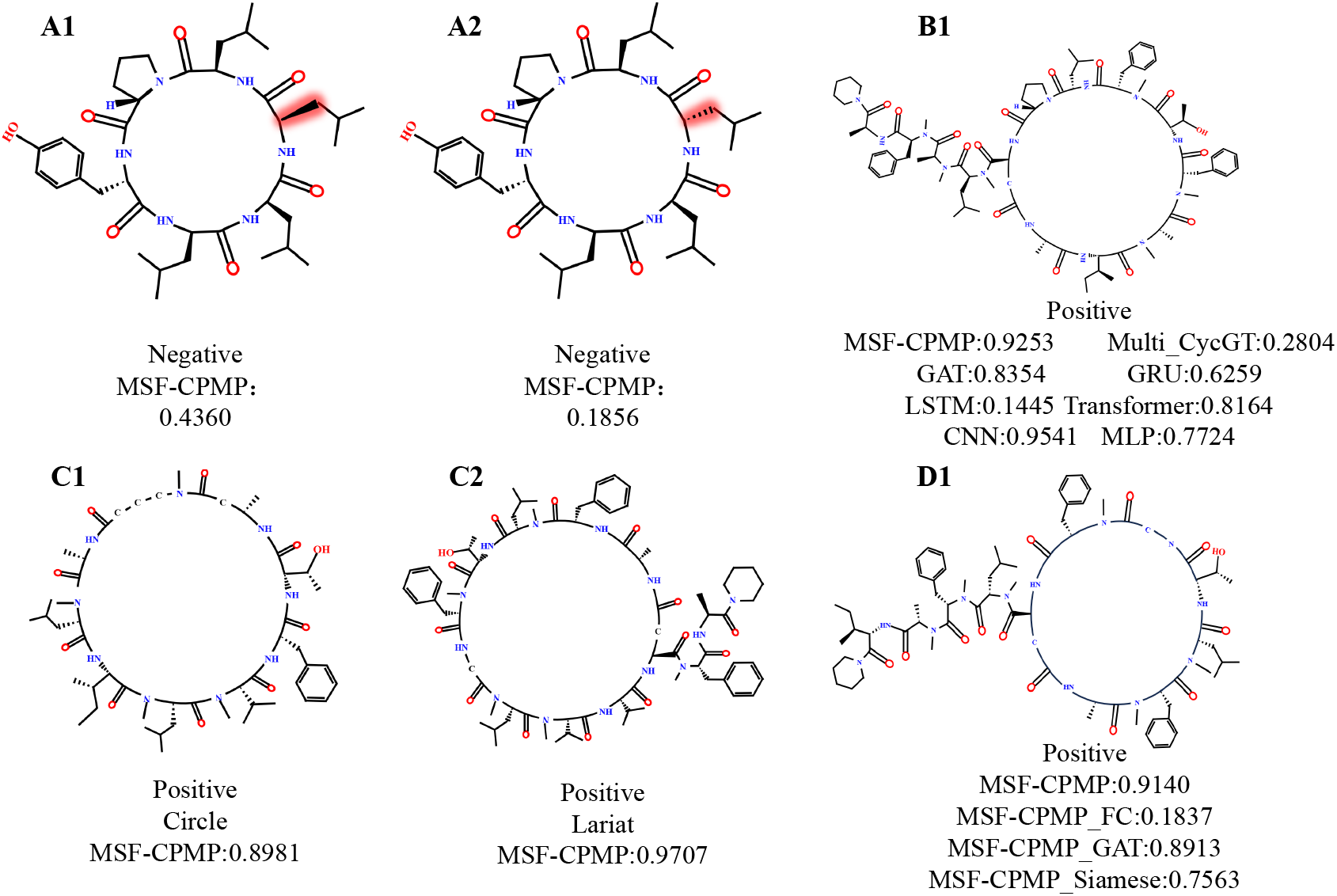
Probability predictions for different categories of cyclic peptides. A1 and A2 depict structurally similar negative cyclic peptides, along with the predicted probabilities from MSF-CPMP for these peptides. B1 represents a positive cyclic peptide. Numbers showing the predicted probabilities from various models for this instance. C1 and C2 illustrate the correct prediction probabilities by MSF-CPMP of cyclic peptides with different cyclization positions. D1 presents the prediction probabilities of cyclic peptides by MSF-CPMP and its various feature combinations. MSF-CPMP FC excludes the physicochemical properties of cyclic peptides during the feature extraction process. MSF-CPMP GAT and MSF-CPMP Siamese exclude graph and SMILES sequence features, respectively, during the feature extraction process. The numbers indicate prediction probability values, with values greater than or equal to 0.5 classified as positive, and values less than 0.5 classified as negative.

The dominance of MSF-CPMP stems from its multi-source feature fusion framework, which synergistically integrates physicochemical descriptors (whose ablation caused an 80.1% performance drop in Fig 4D1), graph features (extracted by GAT to model spatial topology of cyclic scaffolds), and sequence features (processed by Transformer for SMILES semantics). This design enables complementary information fusion, and overcomes limitations of single-modality approaches. Particularly, the incorporation of Transformer and GAT architectures outperforms RNN/GCN in learning long-range dependencies (sequences) and non-local interactions (graphs), while hierarchical feature fusion minimizes critical information loss across deep layers, and thereby comprehensively modelling cyclic peptide permeability properties. Ablation studies validate the importance of each component (Table 3), demonstrating significant performance degradation when individual features are excluded. Overall, MSF-CPMP establishes a reliable, end-to-end framework for cyclic peptide permeability prediction, demonstrating state-of-the-art 0.9546 of ACC across diverse cyclic structures (e.g., Circle/Lariat in Fig 4C1 and C2). Exceptional sensitivity to steric and conformational nuances, and robust multi-modal integration surpasses classical (e.g., LSTM) and specialized (e.g., Multi CycGT) alternatives. Its applicability extends to complex macro-cyclic peptide designs, providing a transformative computational tool for de novo permeable peptide development.

**Table 3.** Ablation testing of MSF-CPMP.

| Model | ACC | Precision | F1_Score | Recall | MCC | TNR | KAPPA | AUC |
| --- | --- | --- | --- | --- | --- | --- | --- | --- |
| **MSF-CPMP** | **0.9062** | **0.8937** | **0.9401** | **0.9916** | **0.7482** | **0.6906** | **0.7269** | **0.9546** |
| MSF-CPMP_FC | 0.8547 | 0.8828 | 0.9007 | 0.9194 | 0.6318 | 0.6606 | 0.6300 | 0.8914 |
| MSF-CPMP_GAT | 0.8234 | 0.8591 | 0.8846 | 0.9116 | 0.5146 | 0.5697 | 0.5106 | 0.8794 |
| MSF-CPMP_Siamese | 0.7937 | 0.8279 | 0.8677 | 0.9116 | 0.4144 | 0.4545 | 0.4045 | 0.8002 |

Three ablated features, 1) MSF-CPMP\_FC signifies the exclusion of physicochemical property features; 2) MSF-CPMP\_GAT indicates the exclusion of molecular graph features, and 3) MSF-CPMP\_Siamese represents the exclusion of SMILES string encoding.

### Ablation experiment

MSF-CPMP aims to predict the CPMP by fusing three sources, graph structural information, SMILES sequences, and physicochemical features. To validate their individual effect, we conducted ablation experiments by ten-fold cross-validation. Three treatment by combinations of these features, including, 1) MSF-CPMP FC, excluded the processing of physicochemical property data from the input features of cyclic peptides. This model incorporated a graph attention network for processing the cyclic peptide graph structural data and a Siamese network for handling the SMILES sequences. 2) MSF-CPMP GAT, omitted the processing of graph structural data for cyclic peptides. Instead, it included a fully connected layer for the physicochemical properties and a Siamese network for processing the SMILES sequences. 3) MSF-CPMP Siamese, removed the Siamese component responsible for processing the SMILES sequence data. This model retained the fully connected layer for physicochemical properties and the graph attention network for handling the cyclic peptide graph structural data.

As shown in Table 3, removing of any feature resulted in decrease in the metric scores, pointing out that all three features make contribution to prediction of CPMP. MSF-CPMP Siamese exhibits much larger decrease as compared to other features (Table 3, Figure S5). This observation implies critical rolea of the Siamese network in processing complex SMILES sequences. It is possible that Siamese network can focus on the position and contextual relationships of each element within the sequence. This also suggests that any combination of two cyclic peptide features affects predictive capability for positive samples. Combining the SMILES sequence (one-dimensional feature) and graph structure (two-dimensional feature) may allow the model to capture features from different spatial perspectives, thereby improving its performance.

The parameters quantity and training time under different feature combinations were analyzed to investigate the impact of multi-source features on model complexity. The parameter size in the MSF-CPMP model was approximately 17.98 MB (Table S7). After removing the Siamese network (MSF-CPMP Siamese), the model parameters decreased more than half of training time, indicating that the Siamese network had a significant impact on prediction of CPMP, which was consistent with the previous data [21]. Additionally, the model was more complex than Multi CycGT, with parameter size exceeding Multi CycGT by approximately 8.75 MB, which explains the longer training time required for the model. The synergistic effect of the Siamese module on model performance was evaluated by testing MSF-CPMP with different output dimensions, with comparative results detailed in Tables S8 and S9.

### Effect of cyclization position of cyclic peptides

CycPeptMPDB includes two types of cyclic peptide molecular shapes, Circle and Lariat. Peptides with cyclization at both the N-terminus and C-terminus are categorized as Circle peptides, while cyclization that occurs not at the sequence ends, but rather between the peptide ‘s terminal and side chains, is classified as lariat. Specifically, such cyclized structures are categorized based on their unique characteristics. Fig 5 displays the distribution ratio of cyclization types and a larger proportion of Circle peptides than Lariat ones. We divided the datasets into Circle and Lariat categories. The result of ten-fold cross-validation (Table 4 and Fig 5) found that the number of Lariat peptides tested was smaller, but the average metrics for ACC, precision, F1 score, AUROC, under cross-validation were higher than those for Circle peptides. Noted is that the Recall values for Lariat and Circle peptides were similar; however, the average TNR for Lariat peptides surpassed that of Circle peptides by 0.2965. This indicates that the MSF-CPMP model is sensitive to both positive and negative samples of Lariat-type peptides, demonstrating superior predictive capability.

**Table 4.** Effect of different cyclic position of cyclic peptides on performance.

|  | ACC | Precision | F1_Score | Recall | MCC | TNR | KAPPA | AUC |
| --- | --- | --- | --- | --- | --- | --- | --- | --- |
| Circle | 0.8674 | 0.8712 | 0.9220 | 0.9805 | 0.5263 | 0.4171 | 0.4838 | 0.9463 |
| Lariat | 0.9085 | 0.9050 | 0.9400 | 0.9781 | 0.7570 | 0.7136 | 0.7446 | 0.9830 |

**Fig 5.**
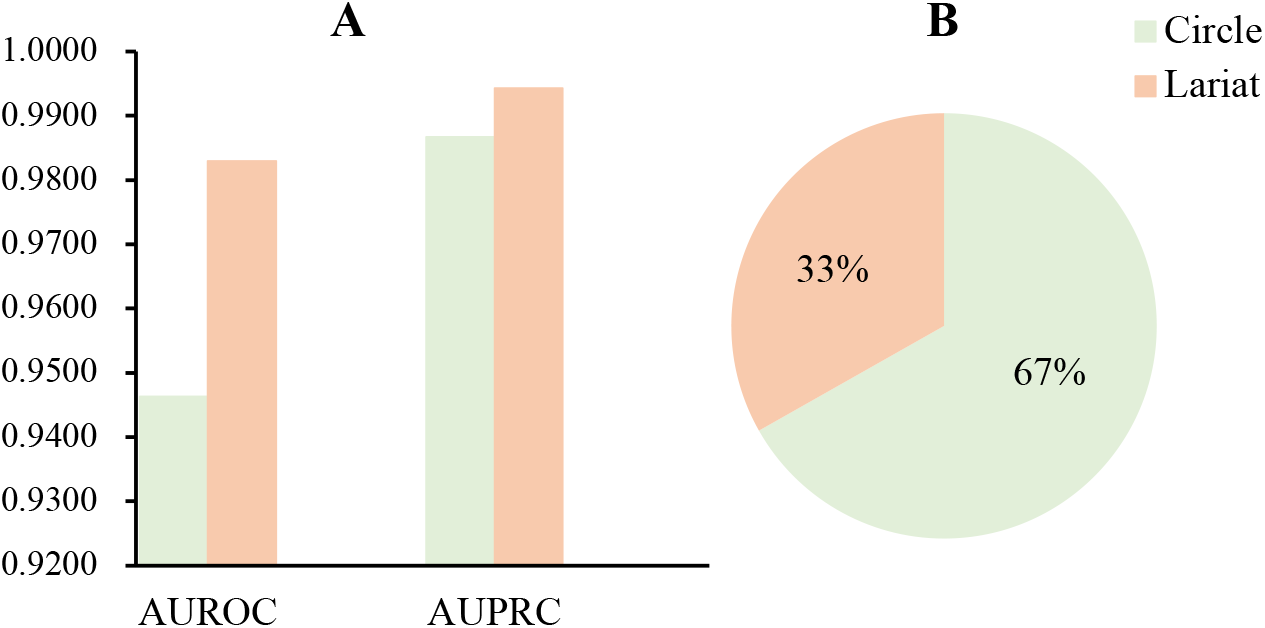
Performance and distribution of Circle and Lariat cyclization positions. (A) Comparison of the performance of Circle and Lariat cyclization positions, including AUROC and AUPRC metrics. (B) The distribution ratios of these cyclization positions.

### Distribution of MSF-CPMP features

To investigate effect of the MSF-CPMP features from different data sources, we used t-SNE to visualize the embedded features before training. After training, the embedded features from the last multi-source feature fusion module were extracted and mapped into a two-dimensional clustering. As shown in Fig 6, the data before training appeared randomly distributed, but after training, the positive and negative interfaces were clearly separated in the embedding space. This result indicates classification features for validating prediction of CPMP using the MSF-CPMP model.

**Fig 6.**
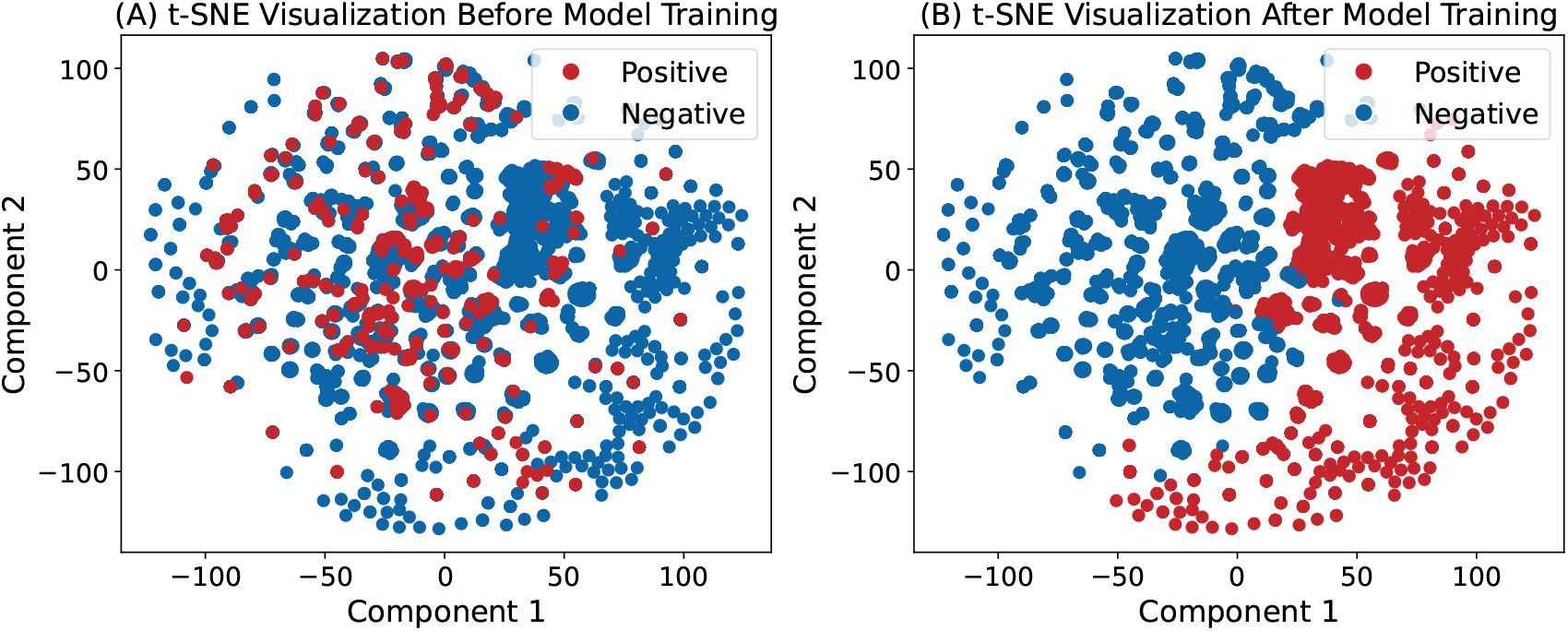
Feature Distribution using t-SNE. (A) distribution of features before model training. (B) distribution of features after model training.

### Effect of different cyclic peptide datasets

To validate the generalization performance of the MSF-CPMP model, three additional cyclic peptide datasets from the CycPeptMPDB were utilized, Caco2, MDCK, and RRCK. Each dataset contains positive and negative samples. The differences are due to experimental methods and cell lines used for predicting permeability. To ensure the reliability of the experimental data, ten-fold cross-validation was conducted on the three datasets using the same parameters as in CycPeptMPDB PAMPA dataset. It is noteworthy that, due to the limited data volume, different batches were employed for training. For the Caco2 dataset, 128 batches were tested, while the other two datasets utilized 8 batches each. The results in Table 5 and Figure S6 found that data volume did not present a significant advantage. Particularly on the MDCK dataset, the performance was optimal, with a Recall value of 1.0000, achieving complete correct predictions for positive samples. This may be attributed to the more prominent and robust features present in the experimental conditions and cell line of the MDCK data. Overall, the model effectively identified and captured the hidden internal features within the cyclic peptides, demonstrating that MSF-CPMP possesses strong generalization capability.

**Table 5.**
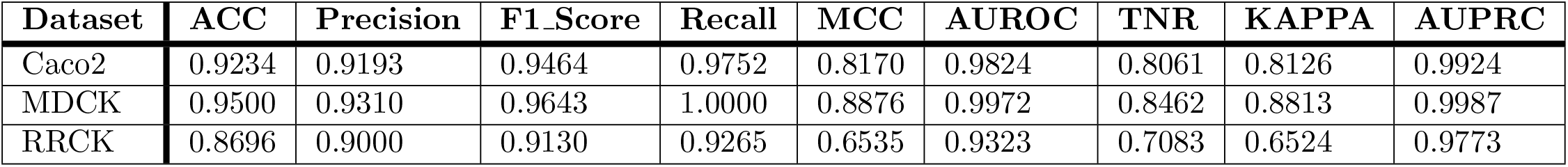
Effect of cyclic peptides measured by different methods Caco2, MDCK, and RRCK on MSF-CPMP performance.

| Dataset | ACC | Precision | F1_Score | Recall | MCC | AUROC | TNR | KAPPA | AUPRC |
| --- | --- | --- | --- | --- | --- | --- | --- | --- | --- |
| Caco2 | 0.9234 | 0.9193 | 0.9464 | 0.9752 | 0.8170 | 0.9824 | 0.8061 | 0.8126 | 0.9924 |
| MDCK | 0.9500 | 0.9310 | 0.9643 | 1.0000 | 0.8876 | 0.9972 | 0.8462 | 0.8813 | 0.9987 |
| RRCK | 0.8696 | 0.9000 | 0.9130 | 0.9265 | 0.6535 | 0.9323 | 0.7083 | 0.6524 | 0.9773 |

## Discussion

In this study, we introduce MSF-CPMP, a novel method for predicting membrane permeability of cyclic peptides through multi-source feature fusion. MSF-CPMP integrates a Siamese network (a parameter-shared Transformer), Graph Attention Mechanism (GAT), and Multi-Layer Perceptron (MLP) to effectively capture SMILES sequences, graph structures, and physicochemical properties of cyclic peptides. The inclusion of residual graph-attention layers is motivated by recent evidence that such architectures outperform conventional graph convolutions across diverse molecular property endpoints [22], while Transformer language models over SMILES strings have been shown to encode long-range chemical dependencies inaccessible to graph-only encoders [23]. By fusing multi-source data via a multi-layer adjacency perceptron, the model achieves precise permeability predictions. Comparative evaluations against other machine learning and deep learning models on benchmarking datasets demonstrate the superiority of MSF-CPMP performance in both classification and regression metrics.

Traditional learners trained solely on handcrafted descriptors plateau below AUROC 0.85 [24], whereas MSF-CPMP achieves AUROC 0.9546, surpassing recent MAT-based CPMP (0.901) [17] and CycPeptMP (0.887) [16] by 5.3–6.7 points using the same dataset from CycPeptMPDB PAMPA. Also, MSF-CPMP can gain lower metrics MSE and MAE, i.e., 0.092 and 0.242, respectively, lower than MAT-based CPMP (0.169 and 0.308), CycPeptMP (0.271 and 0.352), and MultiCycPermea (0.160 and 0.280) [25].

This performance gap underscores how single-modality approaches fail to capture complementary structural-sequence relationships essential for cyclic peptides. Furthermore, the newly developed model reveals the enhanced predictive accuracy for Lariat-type compared to Circle-type of cyclic peptides, while evaluations on three independent datasets (Caco-2, MDCK and RRCK) confirm its robust generalization. The strong performance of the MSF-CPMP model can be attributed to several factors. First is that MSF-CPMP can fuse multi-source features from cyclic peptides, particularly through the introduction of the graph attention mechanism to extract crucial information from the cyclic peptide graph structure. Secondly, the Siamese network utilized in MSF-CPMP captures SMILES representations, addressing the information disparity inherent in cyclic peptides. Moreover, the feature fusion strategy employed by MSF-CPMP significantly enhances prediction of membrane permeability for cyclic peptides. Overall, MSF-CPMP represents a promising approach that leverages deep neural network techniques for predicting the membrane permeability of cyclic peptides.

The exceptional performance of MSF-CPMP arises from its innovative integration of multi-source information, notably through GAT for extracting pivotal graph structural features and the Siamese network for addressing information disparities in SMILES representations. This synergistic feature fusion strategy markedly enhances permeability prediction, positioning MSF-CPMP as a transformative tool in cyclic peptide drug design. When benchmarked against the strongest contemporary deep-learning baselines, such as the MAT-based CPMP, the multi-level fusion network CycPeptMP and MultiCycPermea [25], MSF-CPMP secures absolute AUROC gains of five to seven percentage points. This synergistic feature-fusion strategy markedly enhances permeability prediction, positioning MSF-CPMP as a transformative tool in cyclic-peptide drug design. The t-SNE visualization further underscores the model ‘s ability to discern discriminative features, with distinct separation of permeable and non-permeable peptides, highlighting its potential to advance precision in bioactive peptide development.

Despite these achievements, there are still limitations including all physicochemical properties without identifying key influencers of permeability hampers interpretability. Notably, the model correctly prioritises permeability-enhancing motifs such as intramolecular hydrogen bonds and backbone N-methylation, both of which lower desolvation penalties and facilitate passive diffusion [26, 27]. Future efforts should employ feature selection techniques to isolate critical descriptors, thereby enhancing model transparency. Additionally, while MSF-CPMP excels across diverse datasets, validation in real-world drug screening contexts is essential to confirm its practical utility and resilience.

To further elevate performance, integrating pre-trained language models like ESM-2 or ProtBERT/ProtT5 for SMILES processing could enrich contextual representations of cyclic peptides. Moreover, we will incorporate fast 3D conformer generation and active learning for underrepresented stereochemistry, building on emerging GNN-Transformer hybrids that benefit from iterative data expansion [28, 29], which was shown to boost generalization by 11-14% in comparable architectures, suggesting a promising direction for expanding MSF-CPMP ‘s chemical-space coverage. Expanding the framework to incorporate additional medical data, chemical descriptors, or applications to broader biological systems, such as blood-brain barrier permeability, would amplify its versatility in drug discovery. Lastly, optimizing computational efficiency through model pruning or quantization could facilitate scalability, enabling seamless integration into high-throughput drug screening pipelines.

In summary, our deep learning-based MSF-CPMP model performs multi-source feature fusion extracted from SMILES sequences, graph-based molecular structures, and physicochemical properties of cyclic peptides. Comparative evaluations demonstrate the superior performance of MSF-CPMP over existing methods in predicting CPMP, establishing it as an advancement in data-driven approaches that would guide multi-dimensional predictive modeling, particularly as it outperforms descriptor-based models by more than 10% in AUROC [24] and exceeds specialized deep learning frameworks in conformational sensitivity [19]. Furthermore, this model provides a base for tackling complex human diseases, offering significant contributions to computational biology and clinical treatments.

## Material and Methods

### Datasets

CycPeptMPDB http://cycpeptmpdb.com/ is the largest web-accessible database of membrane permeability of cyclic peptides. This database was established by Tokyo Institute of Technology and collects 6,941 cyclic peptide entries measured using four different methods, namely, PAMPA, Caco-2, MDCK, and RRCK [30]. The PAMPA dataset comprises 6,941 cyclic peptide entries. The Caco-2 dataset contains 649 cyclic peptide entries with human colorectal carcinoma cells as the subject. MDCK (Madin-Darby Canine Kidney) and RRCK (Rat Proximal Cell Kidney) datasets, employing canine kidney cells and rat cortical kidney cells, respectively, contain 448 and 28 entries. This database incorporates detailed molecular information including SMILES notation, computationally generated 3-dimensional structures, and physicochemical properties. We used this database to create our input data of permeability assessment. In data pre-processing, physicochemical properties with predominant values (greater than 99%) were eliminated.

Based on the distribution of permeability provided from CycPeptMPDB (Figure S1), a threshold value of log Pe was used to distinguish between permeable and non-permeable peptides. Values above -6 were classified as positive samples, that generates a positive-to-negative ratio of 61.36%. In comparison, the negative samples in CycPeptMPDB are 38.64%, accounting for 89% of cyclic peptide number, Caco2 for 8%, and MDCK and RRCK contributing 1% and 2% respectively.

### Graph Structure-Based Representation Using Graph Attention Networks (GAT)

The SMILES representation of cyclic peptide chemical structures can be converted into molecular graphs using RDKit [31]. In these graphs, nodes represent atoms and chemical atoms, while edges represent chemical bonds connecting these atoms [32]. Formally, we define a cyclic peptide as a graph *G* = (*V, E*), where *V* denotes the node set and *E* denotes the edge set, with a feature vector **h**_*i*_ derived from encoding fundamental atomic properties, which serves as the feature vector for representing the state at each node.

The core algorithm of GAT updates node features through a self-attention mechanism [33]. Given a set of neighbouring nodes, the node ‘s feature comparison information comprises the following steps:

#### (1) Attention Coefficient Computation

For each pair of adjacent nodes (*i, j*), their attention coefficient is calculated as:

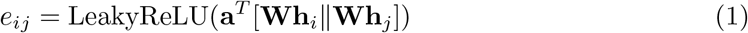

We represent **W** learnable weight matrix, **a** denotes another learnable weight vector, ∥ indicates concatenation operation, LeakyReLU serves as the activation function, and *T* denotes matrix transposition.

#### (2) Attention Normalization

The attention coefficients are normalized to obtain final attention weights:

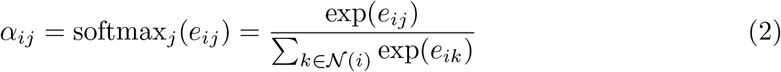

*j* represents all neighbouring nodes of node *i*.

#### (3) Node Representation Update

Finally, the feature vector representation of target node *i* is updated through a weighted sum of neighbouring node features using attention weights:

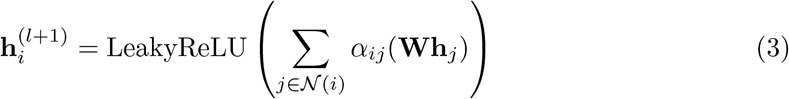

### SMILES-Based Representation Using Siamese Network

The SMILES sequence of cyclic peptide encodes chemical information [34], effectively converting molecular structures into analysable linear strings. To ensure distinct and complete representations of each cyclic peptide SMILES sequence, we employ tokenization to eliminate lexical ambiguities. The detailed tokenization steps are demonstrated in Figure S3, which embeddings to ensure uniform distribution in high-dimensional space, following these computational steps:

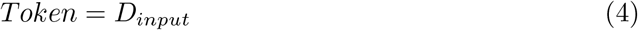

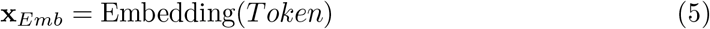

where *D*_*input*_ represents the input token dictionary and **x**_*Emb*_ denotes the embedding vector.

Subsequently, these SMILES embedding vectors **x**_*Emb*_ was processed through two identical Transformer encoders with shared parameters [35, 36]. Recognizing the significance of positional information within SMILES sequences, we implemented positional encoding using a combination of sine and cosine functions:

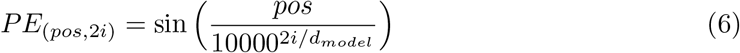

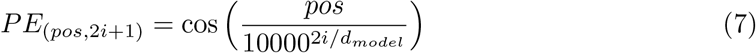

where *PE* denotes positional encoding, *pos* represents the character ‘s position in the input sequence, *i* indicates the encoding vector dimension, and *d*_*model*_ represents the model ‘s embedding dimension.

The combined positional encoding and embedding vectors serve as input **x** to the Transformer encoder, as illustrated in Figure 1A. The Transformer architecture is based on two key components: self-attention mechanism and feed-forward neural networks.

#### (1) Self-Attention Mechanism

The self-attention mechanism captures comprehensive semantic features by analyzing short-term and long-term sequential relationships within SMILES sequences. It begins by performing a linear transformation on the input vectors to derive query, key and value vectors through scaled dot-product attention. Following this, the attention scores are normalized to produce attention weights. Finally, these attention weights are applied to perform a weighted sum of the value vectors to produce self-attention output. The process involves:

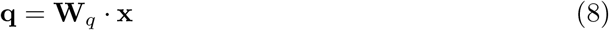

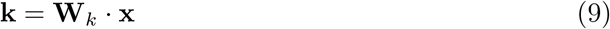

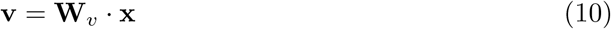

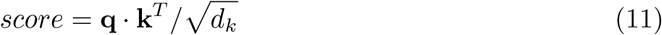

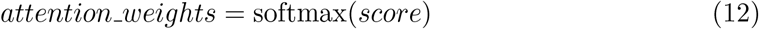

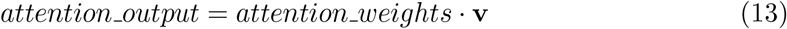

where **x** represents the input vector, **W**_*q*_, **W**_*k*_, and **W**_*v*_ are learnable weight matrices for **q, k** and **v** denote query, key, and value vectors respectively. *score* is attention score, *T* indicates matrix transposition, *d*_*k*_ represents the dimension of **k**. *softmax* serves as the normalization activation function.

#### (2) Feed-Forward Neural Network

Following the attention layer, a fully connected network is implemented with residual connections to the attention layer. Through non-linear processing and feature space expansion, this significantly enhances the model ‘s capability to capture SMILES sequence characteristics and properties.

The dual Transformer architecture with shared weights enables global modelling of SMILES sequence dependencies, unrestricted by sequence length and effectively capturing long-range dependencies. The outputs undergo max-pooling to reduce sequence length while preserving crucial information. Finally, Hadamard product computation addresses feature disparities between sequences and enhances feature representation quality.

### MLP-based Physicochemical Feature Extraction

The physicochemical properties of cyclic peptides demonstrate significant influence on downstream task prediction and serve as crucial inputs to the MSF-CPMP model. Given the high dimensionality of cyclic peptide features, a two-layer fully connected architecture was implemented for feature embedding. Complex features beneficial to the prediction task were learned through this structure, followed by batch normalization to accelerate training and enhance learning capability. This process significantly provides more accurate and robust feature representations for the subsequent feature fusion module.

### Multi-Source Feature Fusion Based on Multi-Layer Perceptron

In the domain of multi-feature fusion tasks, challenges such as excessive parameter counts, computational latency, and high model complexity are particularly prominent. To address these challenges, a two-layer MLP architecture was adopted as the fundamental structure of the feature fusion module in MSF-CPMP. This implementation enables accurate and rapid prediction of cyclic peptide permeability.

### Models selected for comparative analysis

To benchmark MSF-CPMP performance, eight traditional machine learning models were selected. They are: Categorical Boosting Machine (CatBoost) [37], which optimizes categorical features; Light Gradient Boosting Machine (LGBM) [38], which employs histogram-optimized gradient boosting; Random Forest (RF) [39], which builds a decision tree ensemble; k-Nearest Neighbours (KNN) [40], which classifies via nearest neighbours; Support Vector Machine (SVM) [41], which separates classes with a hyperplane using polynomial and radial basis function kernels; Decision Tree (DT) [42], and XGBoost [43]. In the deep learning category, MSF-CPMP is a deep learning-based model. Thus we also selected ten deep learning models. They include Multi-modal Cyclic Graph Transformer (Multi CycGT) [16], Convolutional Neural Network (CNN) [44], Graph Convolutional Network (GCN) [45], Multi-Head Attention (MHA) [35], Recurrent Neural Network (RNN) [46], Long Short-Term Memory (LSTM) [47], and Gated Recurrent Unit (GRU) [48], as well as standard Transformer and Multi-Layer Perceptron (MLP) architectures. Among these models, Multi CycGT represents a multimodal architecture designed for cyclic peptide permeability prediction. It integrates graph-structured inputs via GCN, SMILES sequences via a Transformer, and physicochemical descriptors via fully connected layers.

### t-Distributed Stochastic Neighbor Embedding (t-SNE)

t-SNE is implemented to transform high-dimensional data into lower dimensions, facilitating the observation and analysis of internal data structures [49]. Through this optimization process, low-dimensional points effectively reflect the structural relationships present in the high-dimensional space. This enables visualization and examination of the learned feature embedding distribution characteristics.

### Evaluation of Models through ten-fold cross-validation

Similarly, we examined peptide models through ten-fold cross-validation. The input datasets are partitioned into training, testing, and validation sets in ratio of 8:1:1. This approach effectively mitigates data partition bias through averaged measurements and standard deviations. Each data also ensures representation of both positive and negative classes. The permeasbility of the positive samples is predicted correctly and gives label “1”. By contrast, in the negative samples, the permeability is predicted incorrectly and gives label “0”.

This method ensures that the training and testing datasets were representative of the entire dataset. To increase the reliability of model validation. According to the primary and unbalanced datasets, we carried out the evaluations through two types of metrics, classification and regression.

A total of seven metrics were used in classification. Higher values indicate relatively better performance for the model. We first can calculate TPR (true positive rate), FPR (false positive rate), TNR (true negative rate) and FNR (false negative rate), and the use of ROC (area under receiver operating characteristic curve), AUPRC (area under precision-recall curve), ACC (Accuracy), Precision, F1 Score, Recall, MCC (Matthews Correlation Coefficient), to make comprehensive evaluation for performance of the model. The mathematical formulations are as follows:

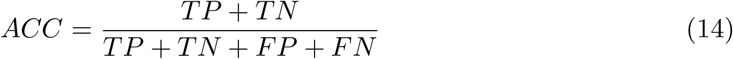

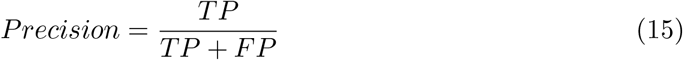

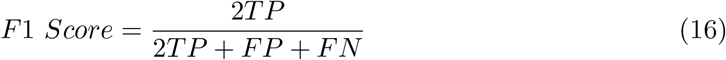

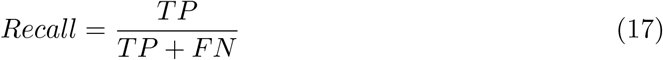

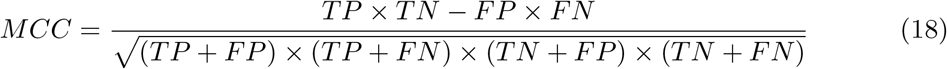

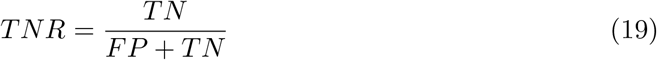

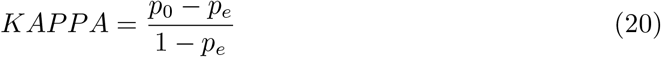

where True Positive (TP) refers to correctly identified positive cases, False Positive (FP) to incorrectly identified positive cases. Conversely, False Negative (FN) denotes incorrectly identified negative cases, and True Negative (TN) represents correctly identified negative cases. For the KAPPA coefficient calculations *p*_0_ represents the observed agreement between model and actual labels, while *p*_*e*_ is the proportion of cases where predictions align with true values, and *p*_*e*_ denotes the expected agreement by chance between predictions and actual labels, derived from the marginal probability distributions.

Six regression metrics, Mean Squared Error (MSE), Mean Absolute Error (MAE), Root Mean Squared Error (RMSE), and Mean Absolute Percentage Error (MAPE), Coefficient of Determination (*R*^2^) and Pearson correlation coefficient (*r*) are calculated. Smaller values of the first four metrics represent better model performance. Their calculations are defined as follows:

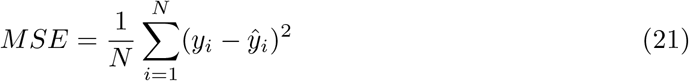

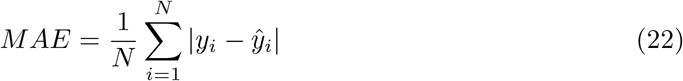

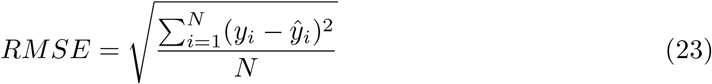

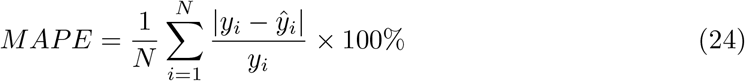

where *N* indicate number of samples, *y*_*i*_ represents the true value of *i*-th sample, while ŷ_*i*_ represents the predicted value of the *i*-th sample.

## Abbreviations

MSF-CPMP: Multi-Source-Fusion Cyclic Peptide Membrane-Permeability Predictor
CatBoost: Categorical Boosting
DT: Decision Tree
KNN: k-Nearest Neighbors
LGBM: Light Gradient Boosting Machine RF Random Forest
SVM: (poly) Support Vector Machine with polynomial kernel
SVM (rbf): Support Vector Machine with radial basis function kernel XGBoost Extreme Gradient Boosting
CNN: Convolutional Neural Network
GCN: Graph Convolutional Network
MHA: Multi-head Attention
RNN: Recurrent Neural Network
LSTM: Long Short-Term Memory
GRU: Gated Recurrent Unit
Multi CycGT: Multi-modal Cyclic Graph Transformer
AUROC: Area Under the Receiver Operating Characteristic Curve
AUPRC: Area Under the Precision-Recall Curve

## Acknowledgments

We thank HKU ITS and GXUN for the computing facilities (HPC/AI) provided.

## Data and Software Availability Statement

The datasets used in this study, along with the source code for MSF-CPMP, are publicly available at https://github.com/wanglabhku/MSF-CPMP.

## Supporting Information

The Supplementary Tables and Figures are available free of charge via the Internet at http://pubs.acs.org.

**Table S1.** Characterization dimensions and parameters of cyclic peptides in MSF-CPMP.

**Table S2.** Information of top ten predicted probabilities of cyclic peptides by MSF-CPMP.

**Table S3.** Performance comparison of MSF-CPMP with machine learning models using classification metrics.

**Table S4.** Performance comparison of MSF-CPMP with machine learning models using regression metrics.

**Table S5.** Performance comparison of MSF-CPMP with deep learning models using classification metrics.

**Table S6.** Performance comparison of MSF-CPMP with deep learning models using regression metrics.

**Table S7.** Number of parameters and training time for different architectures of models.

**Table S8.** Synergistic effect of the Siamese modules on MSF-CPMP performance.

**Table S9.** Results of MSF-CPMP with different output dimensions of modules. D indicates the output dimensions.

**Figure S1.** Distribution of permeability values of cyclic peptides in the CycPeptMPDB database.

**Figure S2.** Proportional distribution of cyclic peptides in the CycPeptMPDB database by different detection methods.

**Figure S3.** The lexical metathesis process for the SMILES sequence.

**Figure S4.** Example of SMILES sequence tokenization.

**Figure S5.** Effects of ablation of three features fused by MSF-CPMP.

**Figure S6.** Effect of cyclic peptides measured by different methods, Caco2, MDCK, and RRCK on MSF-CPMP performance.

